# A Machine Learning approach for assessing drug development risk

**DOI:** 10.1101/2020.10.08.331926

**Authors:** Vangelis Vergetis, Gerasimos Liaropoulos, Maria Georganaki, Andreas Dimakakos, Dimitrios Skaltsas, Vassilis G Gorgoulis, Aristotelis Tsirigos

## Abstract

Characterizing drug development risk – the probability that a drug will eventually receive regulatory approval – has been notoriously hard given the complexities of drug biology and clinical trials. This often leads to an inefficient allocation of resources, and an overall reduction in R&D productivity. We propose a Machine Learning (ML) approach that provides a more accurate and unbiased estimate of drug development risk than traditional models.

## INTRODUCTION

Three is little doubt that drug development is a very risky endeavor. It takes more than 10 years, it costs about $2.5B, and less than 10% that start clinical development are eventually approved by regulators ([1], [2], [3]). Within this context, the ability to accurately assess the risk of drug development (known in the industry as PTRS)^1^ of a particular development program^2^ is critical.

First, it can allow drug development companies to prioritize among the different programs in their pipeline, make better resource allocation decisions, and ultimately increase the productivity of R&D efforts and investment [4]. Second, it can provide a broader view of risk and optimize resource allocation, as companies can also estimate the risk associated with external molecules that they can potentially acquire/in-license.

### Current Approach

In broad terms, the current approach that the industry uses to estimate the PTRS of a development program is based on three main inputs: (i) historical benchmarks driven by the current state/Phase of the program and the specific disease, (ii) expert input from Key Opinion Leaders (KOLs), and (iii) statistical analyses performed by the R&D analytics groups within pharma companies and biotechnology companies. The typical approach is to start with an initial estimate based on (i) above, and subsequently adjust that historical benchmark based on qualitative input from KOLs (e.g., depending on beliefs around the specific biology of the drug, the specific safety and efficacy data, etc.), and input from the analytics team of the sponsor company.

To our knowledge, there have only been early efforts to utilize ML in the process described above. Based on our own experience, we see several reasons for this underwhelming use of ML in drug development to-date – both technical, as well as cultural. Within this context, we describe below an approach that is based on (i) a comprehensive dataset of successes and failures of development programs across the industry, and (ii) ML-driven models, instead of the statistical approaches described above.

## METHODS

### Overall data structure

We propose a systematic approach that incorporates more than 100 different factors (features) across five broad areas:

1. *Clinical trial design:* Choice of primary and secondary endpoints, number of arms, inclusion/exclusion criteria, type of comparator, use of biomarkers, number of patients, etc.
2. *Clinical trial outcomes:* The reported outcomes of the drug in previous studies. For example, the published Phase 2 and Phase 1 data for a drug that is currently transitioning to Phase 3.
3. *Drug biology:* Mechanism of action, modality, genomic data, molecular structure, etc.
4. *Regulatory data:* Any designations by regulatory agencies. For example, prior approvals in other indications, breakthrough therapy designation, etc.
5. *Sponsor characteristics:* The level of experience of the company running the development program in the specific disease area, etc.

To be precise, each development program is represented by a set of 117 features, across the 5 categories mentioned above. The target variable then is whether the program has received regulatory approval by the FDA (or not). In addition, one needs to keep track of when the data is released, especially clinical trial outcomes data: improperly incorporating such data without taking into account the release dates could result in models that suffer from forward-looking biases (see more details below).

We also note here that some of the feature categories above are specific to Therapeutic Areas and/or specific indications. For example, within (2) above, oncology trials measure different outcomes (e.g., ORR, OS, PFS, etc.) than trials in Rheumatoid Arthritis (e.g., ACR 20/50/70 criteria, etc.) or Parkinson’s, COPD, etc. And endpoints can vary even within a Therapeutic Area. For example, while most oncology trials measure outcomes like ORR, OS, and PFS, trials in prostate cancer would measure additionally prostate-specific antigen levels, while in many hematological cancers CRR would be monitored^3^. Similarly, the most relevant drug biology features under (3) above can differ from disease area to disease area. As a result, we argue that “one size doesn’t fit all”: While overarching machine learning models can be trained across therapeutic areas, models that are specifically tailored to an indication (e.g., prostate cancer) or a family of indications (e.g., liquid tumors) are likely to perform better.

Finally, we should emphasize the practical challenges and extensive work required to assemble this data: Dozens of data sources need to be accessed, data needs to be harmonized, consistent ontologies (e.g., for genes, targets, drug modalities, pathways, etc.) need to be created. Our experience also suggests that the need for human curation and quality control, even in the era of “big data” and AI, should not be underestimated. In our view, more than 80% of the effort can typically go in data-related tasks described above, leaving about 20% of the overall effort towards training the different machine learning models. Figure 1 below summarizes the overall methodology used for assembling, curating, and preparing the data for use in our Machine Learning models described in the next section.

**Figure 1:**
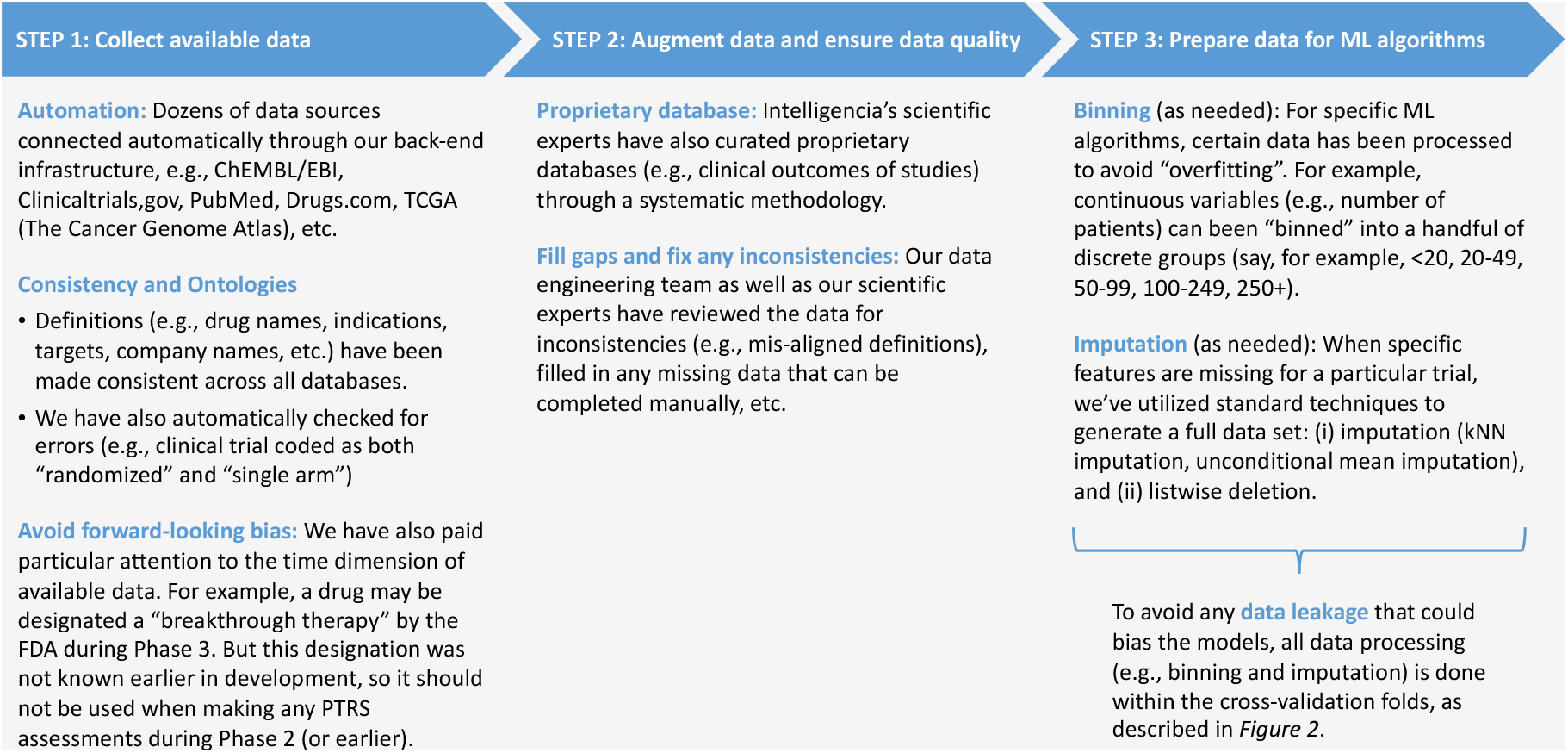
Summary of overall methodology and data curation.

### Prepare data for ML models

While collecting and curating the relevant data is an immense task, there is still work that needs to be done before ML algorithms can be trained. We briefly describe how we use imputation to address gaps in the underlying data, and how specific variables are binned in order to avoid overfitting of the ML models.

Imputation of missing data: Various methods of imputing data were used so as to have as much of a complete dataset as possible, without introducing any bias into our models, as described in more detail below. To better design our data imputation methods, we first analyzed the reason why we would have missing data in the first place. Different data sources have different underlying reasons why they would produce missing data in our dataset. For example, imputing values on the number of arms of a trial should be treated differently to the modality of a drug. Before proceeding on implementing our imputation methods we first:

- Filled in some of the missing data by looking/expanding to additional data sources;
- Made logical assumptions on some missing feature values given other known feature values. For example, a single-group assignment trial cannot be a randomized trial at the same time; and
- Found features that are correlated and could provide information on missing values.

Our imputation methodology steps can be summarized as below:

- For most categorical features, we would fill the missing values of the observations with the most frequent values of that feature in the respective group of observations
- For some continuous features, we would fill the missing values of the observations with the median of that feature in the respective group of observations (i.e., same indication in the same phase of development) if the feature exhibited long tails in its distribution, or with the mean in other occasions.
- For features that we could find strong correlation to other features and that could have a logical explanation, we trained a k-Nearest-Neighbors model to predict the missing values (kNN-imputation) by taking into account the most correlated features of the feature to be predicted.

It is important to note that imputation was done by looking only at the dataset to be used as training, having first completely ring-fenced the test dataset from any data-peeking.

Binning of variables: In order to avoid overfitting our models by allowing the algorithms to break down some predictive continuous features into very small regions with few observations, we implemented a binning methodology whereby we would create a defined number of bins for some continuous features by choosing the optimal thresholds. The choice of the optimal thresholds for the features that underwent binning was left to a univariate algorithm that produced the defined number of bins by maximizing the AUC of the trained dataset.

### Machine Learning algorithms

Once the data described above are collected and curated for all development programs that have been both approved as well as failed in the past ∼20 years, a machine learning model can be trained to identify patterns in all that data and accurately predict the PTRS of the development program of interest.

To be precise, this paper focuses on oncology and our models include clinical trials since 2000. This includes 350 development programs that received regulatory approval, and 4,354 programs that failed to receive regulatory approval. We have also developed similar models in other Therapeutic Areas (e.g., in Inflammation & Immunology, Central Nervous System, etc.) but those are beyond the scope of this paper. Table 1 below summarizes the size of the data set that we used.

**Table 1:** Number of development programs used in this work.

| **Therapeutic Area** | **Number of programs** |  |  |
| --- | --- | --- | --- |
|  | **Successful** | **Unsuccessful** | **Total** |
| Solid tumors | 235 | 3,326 | 3,561 |
| Liquid tumors | 115 | 1,028 | 1,143 |
| **Total Oncology** | **350** | **4,354** | **4,704** |

Within the context described above, we trained several machine learning algorithms: Random forests, k-nearest neighbors, Support Vector Machines (SVMs), logistic regression, and Gradient Boosting Trees.

Below, we articulate some more details around the methodology we used.

#### Forward-looking bia

Most of the features are connected in one way or another to a relevant date (time-dependent) while others are fixed across time. For example, the trial design of a trial is determined by the start of a trial while the mechanism of action of a drug does not change with time. We were very diligent when it comes to collecting and timestamping the data that were used in our models so as not to take into account any kind of information that became known after the date that was used for our predictions and avoid any forward-looking bias in our models. We would therefore look, for example, if and when a particular program received the ‘breakthrough therapy’ designation and only use that information for predictions that encapsulated data after that point in time.

#### ML algorithms selection and evaluation

Nested Cross-Validation was chosen as the method to select, tune and evaluate our Machine Learning models. To avoid any bias that could occur by testing our models in a specific subset of the dataset, we choose to evaluate our models via k-fold validation. Each time we train our models in the k-1 folds and test their performance in the k^th^ fold, k times. We chose to perform 10-fold validation so as to have 90% of the data in each “outer training fold”. The model selection and tuning is done via a process where for each “outer training fold” we run an additional cross-validation where multiple Machine Learning models with a range of possible hyperparameter values are being trained in each “inner training fold” and the ones with the best performance across the “inner testing folds” are being chosen. Figure 2 below illustrates that method. The Area Under the Receiver Operating Characteristic Curve (AUC-ROC) is chosen as the metric to measure the algorithm performance. The models are then trained using the whole “outer training fold” dataset and produce an “out-of-sample” probability for each observation in the test fold dataset. All operations (e.g., binning of variables, data imputation, etc.) are done within each of the cross-validation folds to avoid any data leakage.

**Figure 2:**
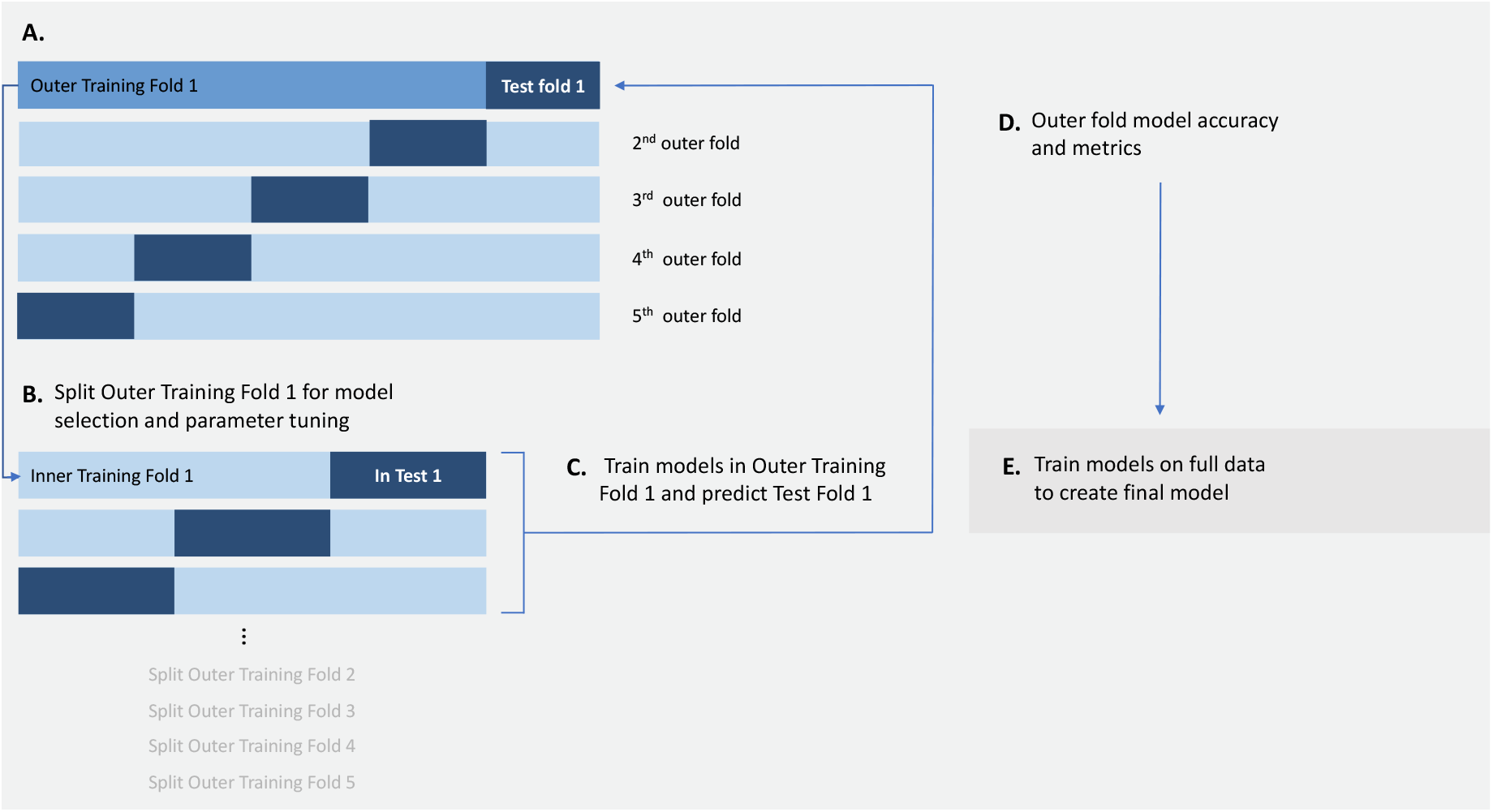
Summary of overall Machine Learning methodology.

#### Overfitting metric

The difference in the AUCs between the training set and the test set for each model across each fold is being saved and monitored as a way to capture any potential overfitting of our models.

#### Prospective analysis

We used prospective analysis to pressure-test the consistency of our models. In the prospective analysis, we create a test set of the known drug development pipelines of the last X years (e.g., drug development pipelines that got approved or failed/discontinued during the last 3 years) and provide as training data only instances that were approved or failed prior to that date. For example, we train a model based on all trials in the 2000-2016 timeframe, and then test the models based on successes and failures that occurred in 2017-2020. Section D below discusses the performance of these models.

Figure 2 provides an overall summary of our ML methodology, and Figure 3 provides some more detail of the training process within each of the nested cross-validation folds.

**Figure 3:**
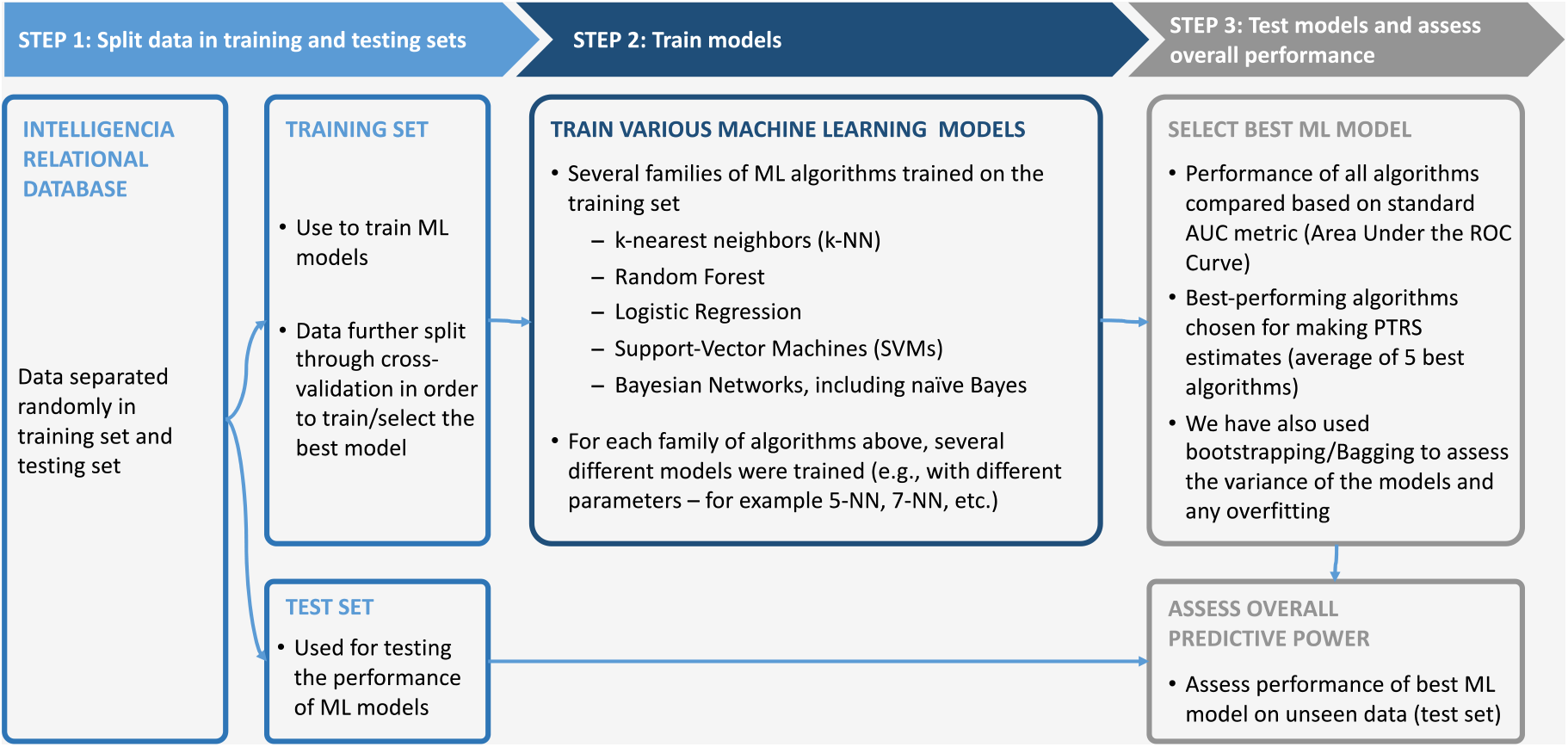
Summary of training and testing of ML models (within each fold)

## RESULTS

### Model performance

The models were rigorously tested under different conditions, for example:

- Different Therapeutic Areas and indications (e.g., Oncology, Inflammation, Immunology, Central Nervous System diseases, etc.). In this paper we focus on Oncology.
- Different stages of development (e.g., Phase 1a/b, Phase 2, Phase 3);
- Different types of molecules (e.g., first-in-class, versus targets that have been previously validated/approved, etc.); and
- Different timeframes (e.g., prior to 2010, 2010-2016, 2017 onwards).

Performance metrics are obviously reported on unseen test sets. Using the standard AUC (area under the ROC^4^ curve) metric for a test set that was not used during the training process, our models achieve a performance of 0.81-0.93 depending on the different scenarios mentioned above. Tables 2-3 below summarize the predictive performance of our algorithms across different scenarios, and Figures 1-16 in the Appendix show the actual ROC curves (and corresponding AUC) for those scenarios.

**Table 2:** Summary of algorithm performance (AUC) in different scenarios.

|  | Phase of program |  |  |  |  |
| --- | --- | --- | --- | --- | --- |
| Example indication(s) | Phase 1 start | Phase 1 end | Phase 2 start | Phase 2 end | Phase 3 start |
| Solid tumors | 0.91 | 0.92 | 0.89 | 0.88 | 0.81 |
| Liquid tumors | 0.91 | 0.91 | 0.92 | 0.93 | 0.81 |

**Table 3:** Summary of algorithm performance (AUC) – prospective testing.

|  | Phase of program |  |  |  |  |
| --- | --- | --- | --- | --- | --- |
| Example indication(s) | Phase 1 start | Phase 1 end | Phase 2 start | Phase 2 end | Phase 3 start |
| Solid tumors | 0.89 | 0.92 | 0.90 | 0.89 | 0.79 |
| Liquid tumors | 0.87 | 0.87 | 0.95 | 0.94 | 0.76 |

To illustrate one specific example, when testing the algorithm across several hundred randomly chosen solid tumor programs at the beginning of Phase 2, the AUC is 0.89 for solid tumors and 0.92 for liquid tumors. Put another way, the algorithm correctly predicts more than 79% of the development programs that eventually receive regulatory approval, and more than 82% of the programs that don’t for solid tumors. For liquid tumors, the algorithm correctly predicts more than 81% of the development programs that eventually receive regulatory approval, and more than 91% of the programs. This performance is intriguing, particularly given how early those predictions are made (development programs at the beginning of Phase 2 can be 4-6 years away from a regulatory decision), and how complex the overall problem is. Further, we note that the 95% confidence intervals for those AUC values are within +/-0.03 from the stated values. In other words, with 95% confidence the AUC of the algorithm for solid tumors at the beginning of Phase 2 is 0.86-0.92, and for liquid tumors at the beginning of Phase 2 is 0.89-0.95.

Perhaps the most important question is whether this model can generalize and, as a result, predict the future – in other words, prospective testing. To explore this scenario, we trained several models using data from development programs that either succeeded or failed in the 2000-2016 timeframe, and then tested the performance of the model on development programs in the 2017-2020 timeframe. Table 3 summarizes those results. To use the same example of early Phase 2 solid tumor programs above, the model still achieved an AUC of 0.90 for solid tumors and 0.95 for liquid tumors. Put differently, the algorithm for solid tumors correctly predicted more than 81% of the development programs that eventually received regulatory approval, and more than 85% of the drugs that didn’t in the unseen post-2017 dataset, while for liquid tumors the respective numbers are 85% and 91%.

With respect to the most predictive features within our ML models, those include the clinical outcomes, some elements of the trial design (e.g., choice of endpoints and number of patients), the mechanism of action and related biological characteristics, as well as whether the FDA has granted a breakthrough designation to the molecule being developed.

### Comparison to simple predictive models

In this short section, we compare the predictive power of the ML models developed in this work with the predictive power of simple statistical models. To do this, for each of the models described in Section D above (i.e., for different Therapeutic Areas, stages of development and timeframes) we also developed simple univariate-based models on the most predictive variables in each case.

Specifically, we first used statistical univariate tests to identify the top predictive variables in each instance. For example, the top three predictive variables (based on univariate statistical tests) for oncology programs at the beginning of Phase 2 were the number of patients of the trial, whether it had received orphan drug designation and a metric reflecting the experience that the leading sponsor has accumulated in that therapeutic area. We then used those variables to develop simple regression-based models to predict the PTRS of development programs based solely on those variables. Using solid tumors as an example, Table 4 below summarizes the performance of those statistical models and compares them to the performance of our ML models. We notice that ML models improve the overall predictive power (AUC) by 0.05-0.20. While Table 4 performs this analysis for solid tumors, we observed similar behavior in liquid tumors, and in other Therapeutic Areas. We also note here that the AUC across all three models is lowest at the start of Phase 3. Models in earlier phases of development pick up several predictive signals and distinguish between the majority of trials that eventually fail, and the few that eventually succeed. At the beginning of Phase 3, after programs have successfully completed Phases 1 and 2, the distinguishing characteristics of successful and unsuccessful programs are more nuanced. More work is needed to further improve the performance of models for development programs in Phase 3.

**Table 4:** Comparison of ML vs. statistical regression models for solid oncology programs.

| *Phase of development* | *Model performance (AUC)* |  |  |
| --- | --- | --- | --- |
|  | *ML model* | *Univariate model* | *Multivariate model* |
| *Start of Phase 1* | *0.91* | *0.71* | *0.77* |
| *End of Phase 1* | *0.92* | *0.83* | *0.87* |
| *Start of Phase 2* | *0.89* | *0.75* | *0.78* |
| *End of Phase 2* | *0.88* | *0.73* | *0.78* |
| *Start of Phase 3* | *0.81* | *0.65* | *0.68* |

### Impact of data quality and completeness

In this section we assess the importance of the underlying data when it comes to the predictive power of the ML models. To do so, we created a separate model that relies on the exact same ML methodology (same algorithms, parameters, etc.) as our original models described in Sections C and D above but has limited training data available. More specifically, we used our original training set, but we deleted any data that has been expertly curated by our team of scientific associates (see Step 2 of Figure 1 above) – this primarily included clinical outcomes data, some data on drug biology, and some data on clinical trial design that could not be automatically pulled from sources liked clinicaltrials.gov. We then compared the performance of those two models: The original model trained on the full set of data; and the model described above, that was trained only on publicly available data that has not been expertly curated. Table 5 below summarizes the results of those models. As one can note from the last column, access to expertly curated data improves the overall AUC by 0.15-0.33. While Table 5 performs this analysis for solid oncology programs, we observed similar behavior in liquid tumors and in other Therapeutic Areas.

**Table 5:** Comparison of performance of ML models in solid tumors, trained on different data sets.

| *Phase of development* | *Model performance (AUC)* |  |
| --- | --- | --- |
|  | *Trained on full data* | *Trained on limited data* |
| *Start of Phase 1* | 0.91 | 0.58 |
| *End of Phase 1* | 0.92 | 0.75 |
| Start of Phase 2 | 0.89 | 0.72 |
| End of Phase 2 | 0.88 | 0.73 |
| Start of Phase 3 | 0.81 | 0.61 |

## CONCLUSION

While this work focused on ML-driven decision support models, we emphasize that we do not advocate that drug development decisions should rely solely on the recommendations of machine learning models. Drug development is an incredibly complex process, and there is still “art” involved in it. But we do argue that drug development experts in Portfolio Review Committees and/or Business Development decision-makers can utilize these machine learning models to calibrate and potentially remove any biases from their decision. These models do not rely on just a subset of successful or unsuccessful development programs. They have uncovered complex patterns across all such programs and can offer insights that will allow drug developers to make more informed decisions and with greater confidence.

## ACKNOWLEDGEMENTS

This work has been supported by funds from the Deutsche Forschungsgemeinschaft (DFG), the European Research Council (ERC), and the Helmholtz Association. This research was partially funded by the National Public Investment Program of the Ministry of Development and Investment of Greece/ General Secretariat for Research and Technology.

## SUPPLEMENTAL INFORMATION

**Supplemental Figure 1:**
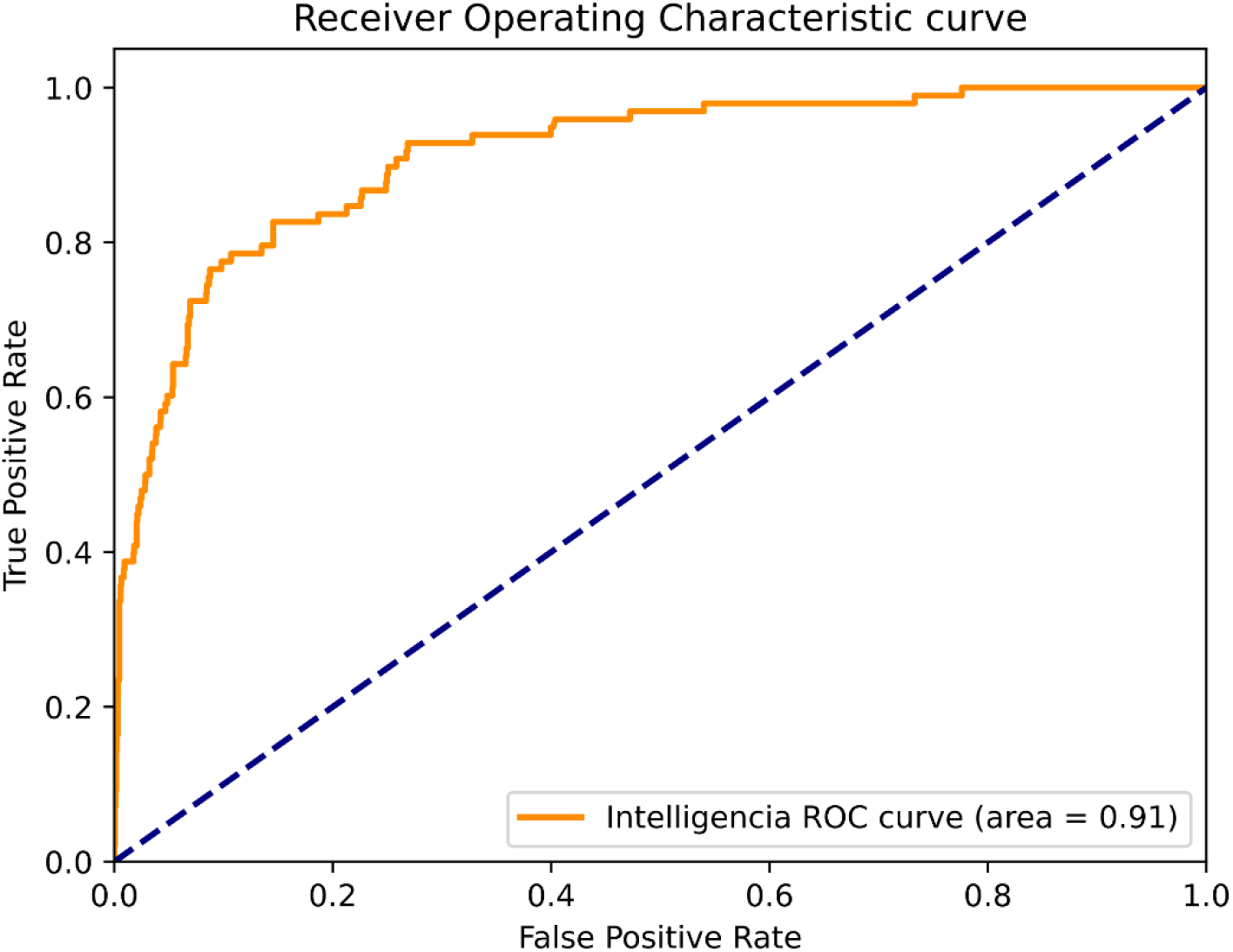
Solid tumors at the beginning of Phase 1.

**Supplemental Figure 2:**
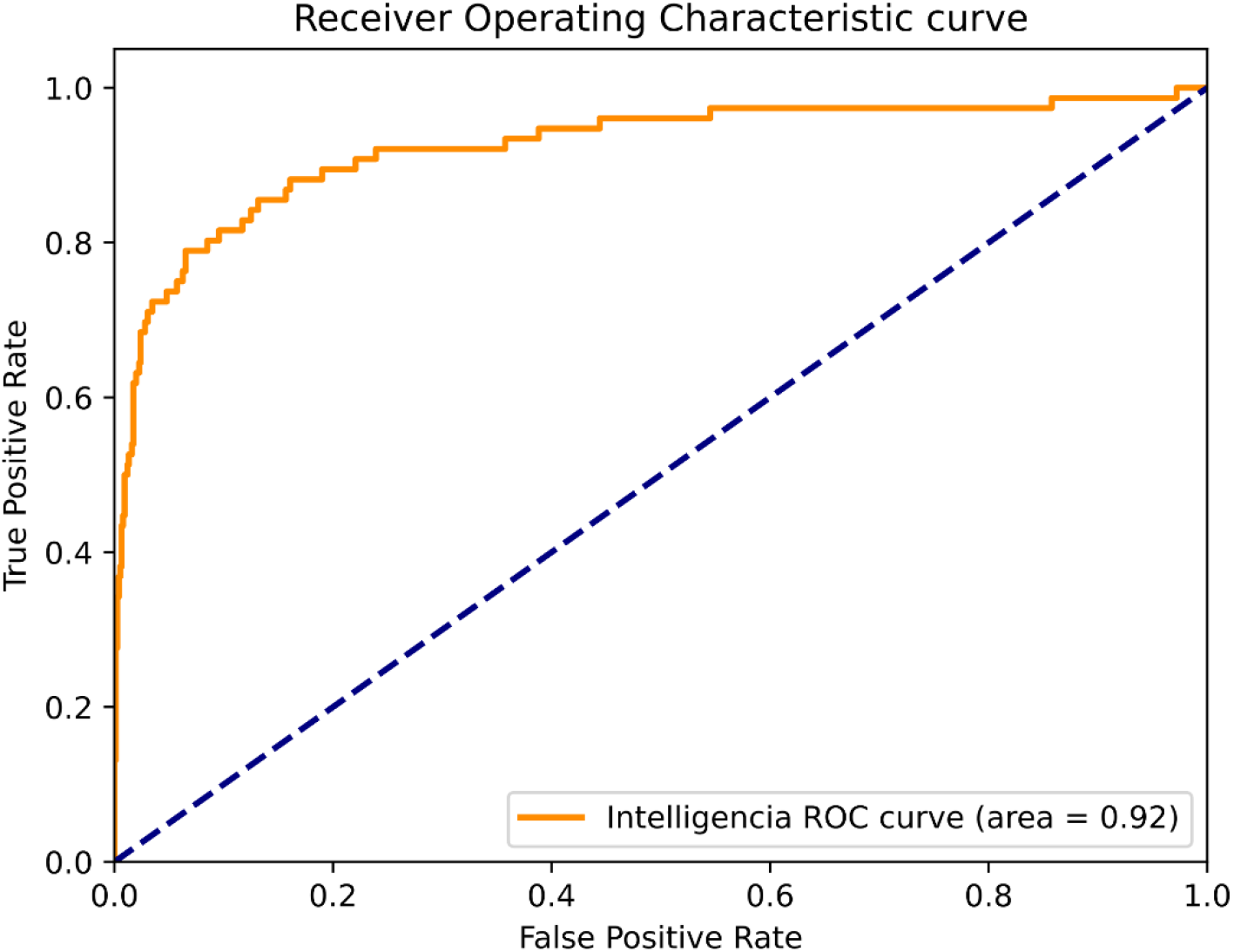
Solid tumors at the end of Phase 1.

**Supplemental Figure 3:**
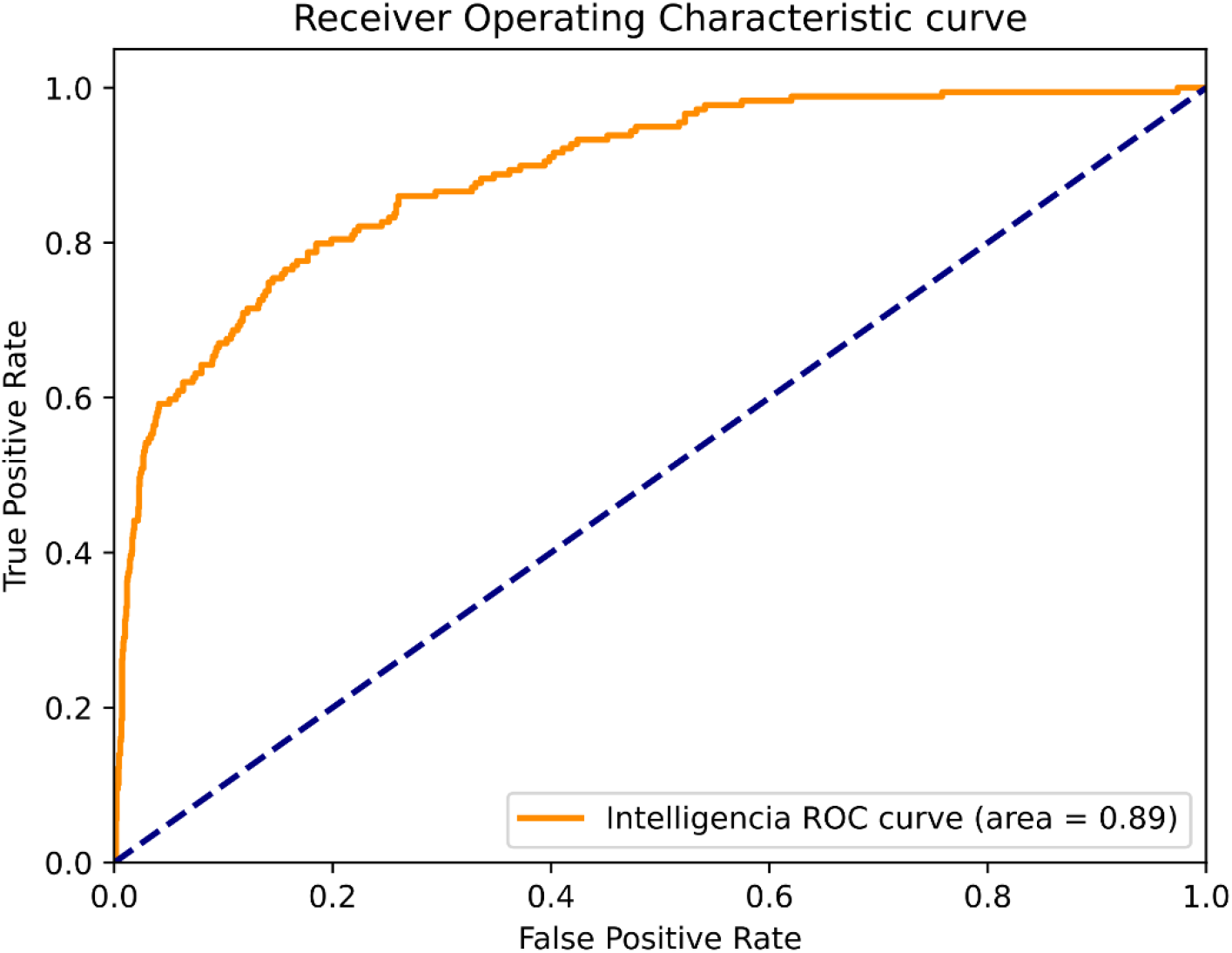
Solid tumors at the beginning of Phase 2.

**Supplemental Figure 4:**
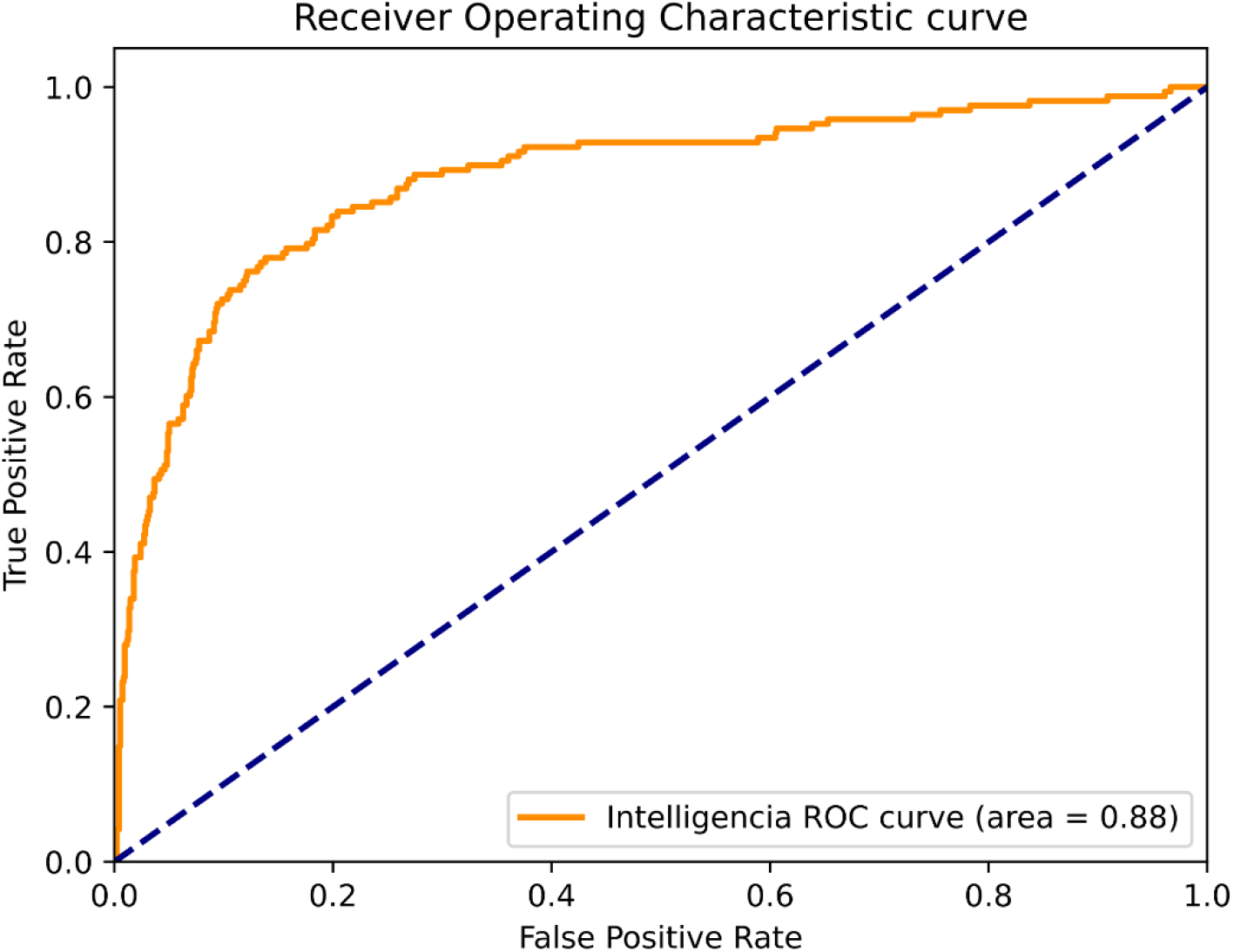
Solid tumors at the end of Phase 2.

**Supplemental Figure 5:**
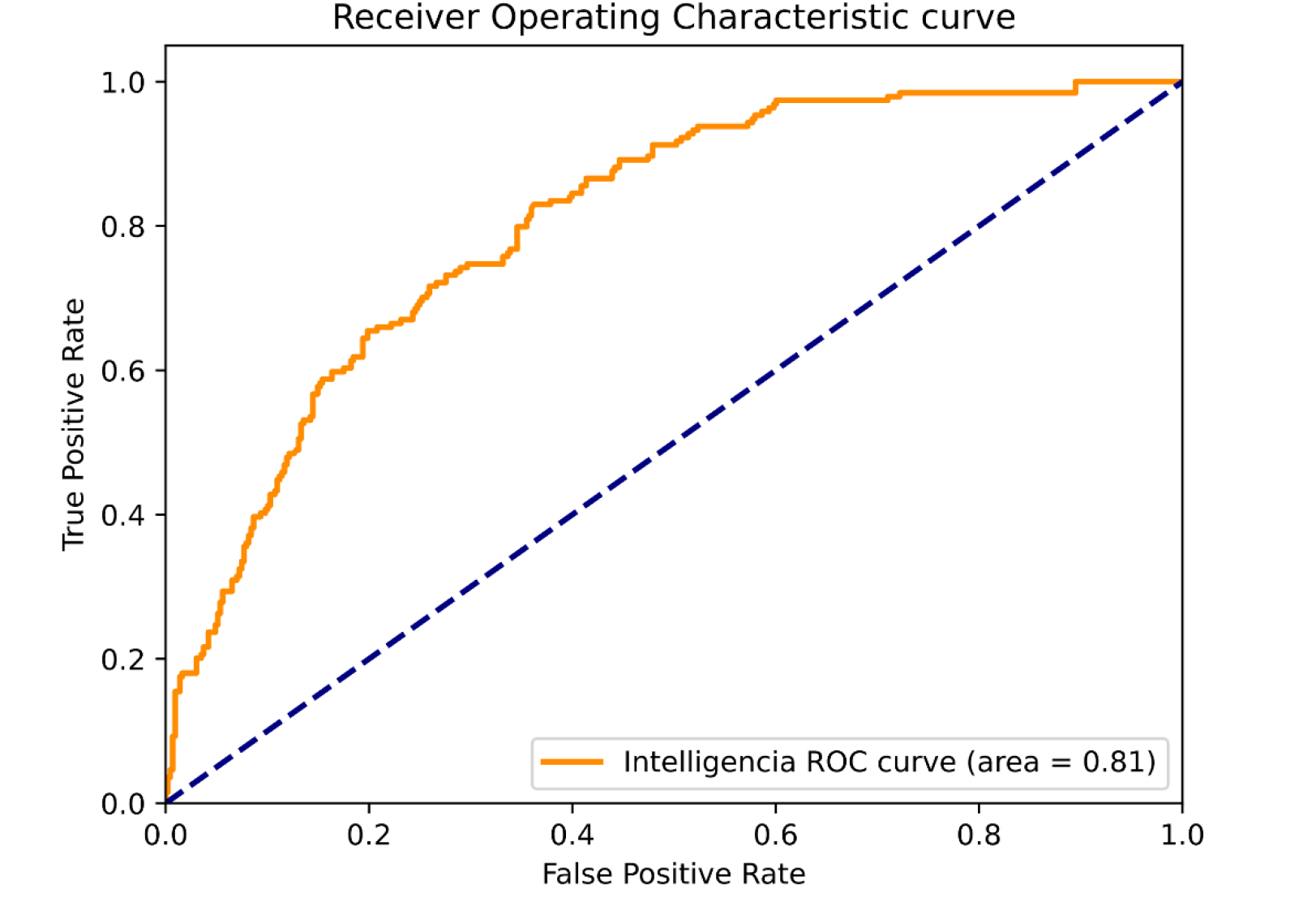
Solid tumors at the beginning of Phase 3.

**Supplemental Figure 6:**
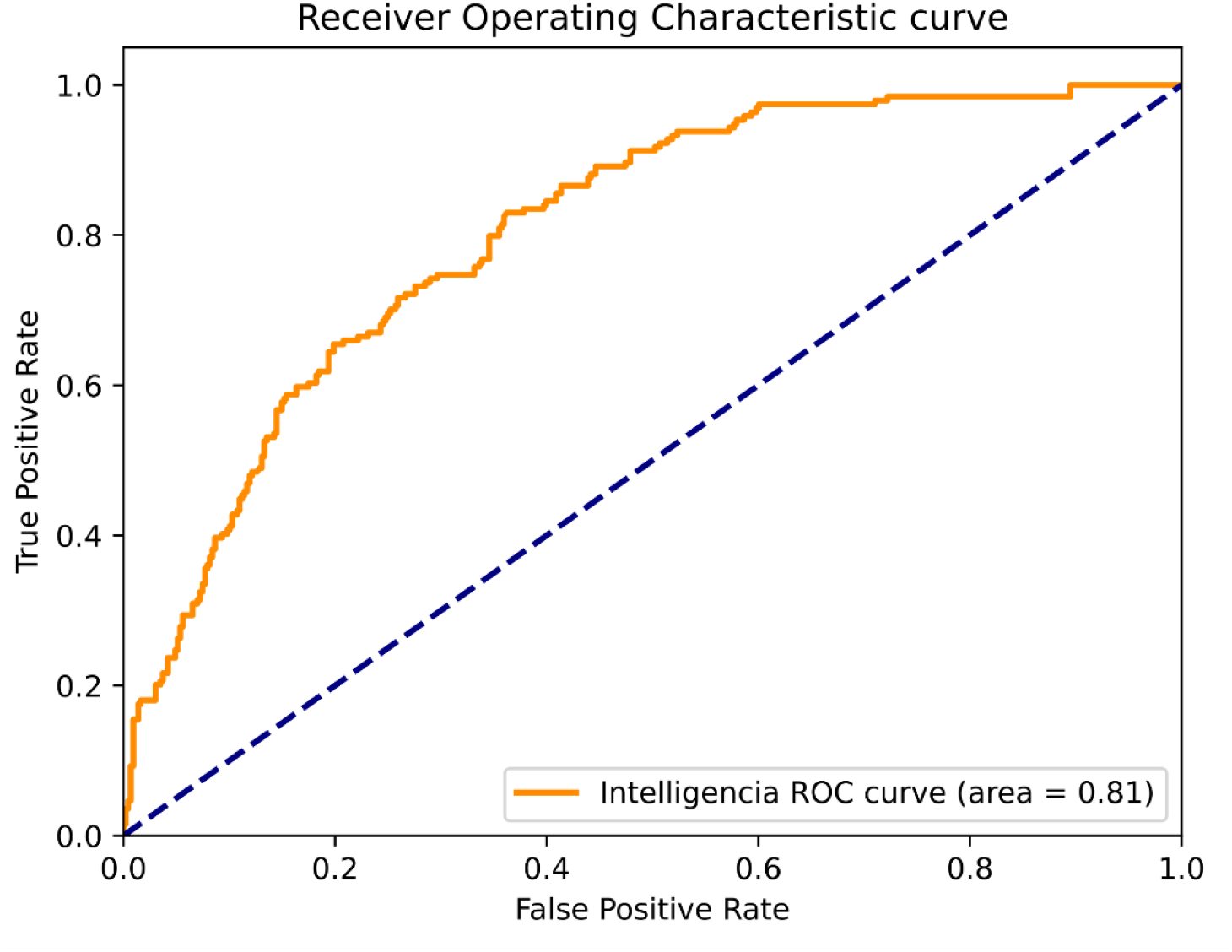
Liquid tumors at the beginning of Phase 1.

**Supplemental Figure 7:**
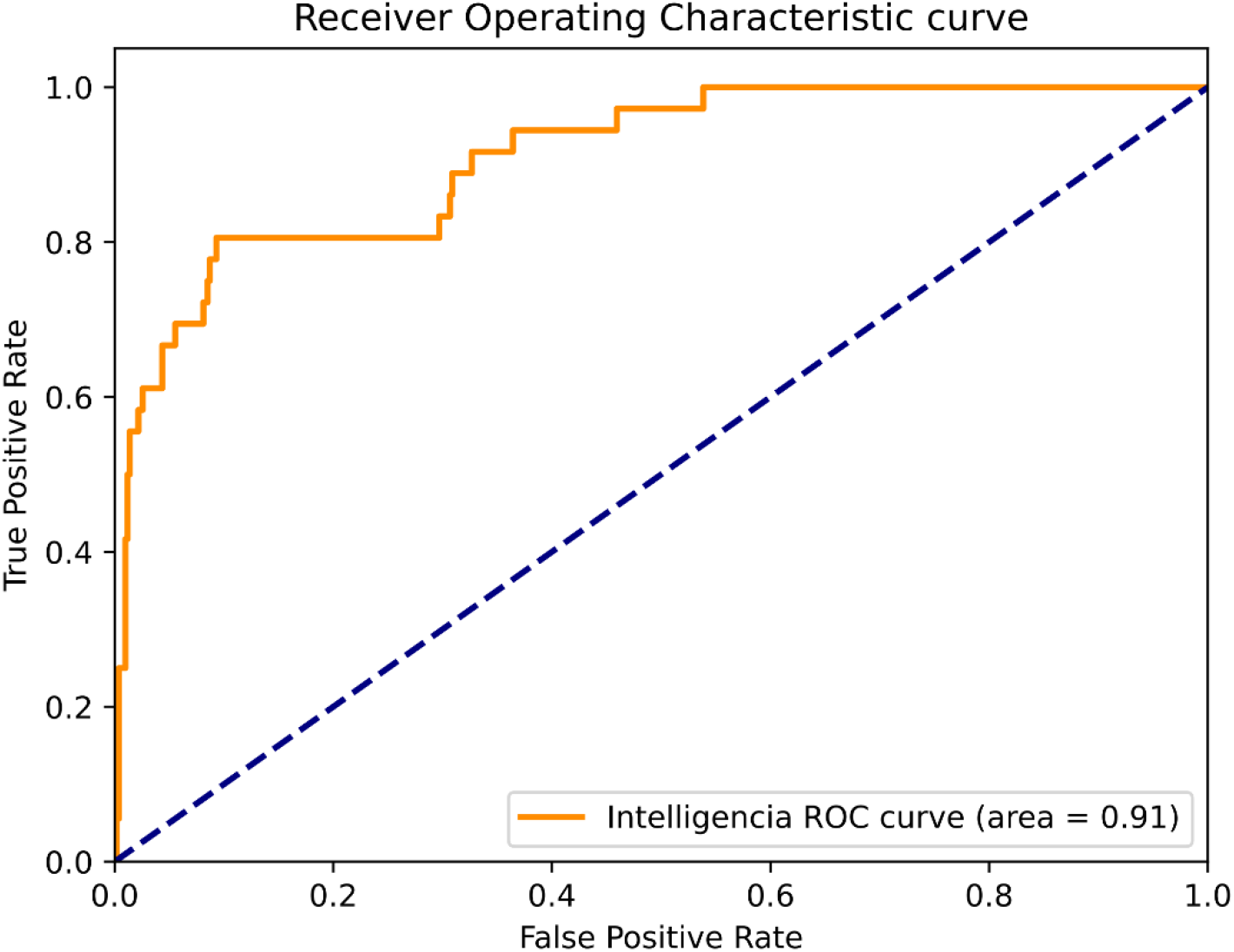
Liquid tumors at the end of Phase 1.

**Supplemental Figure 8:**
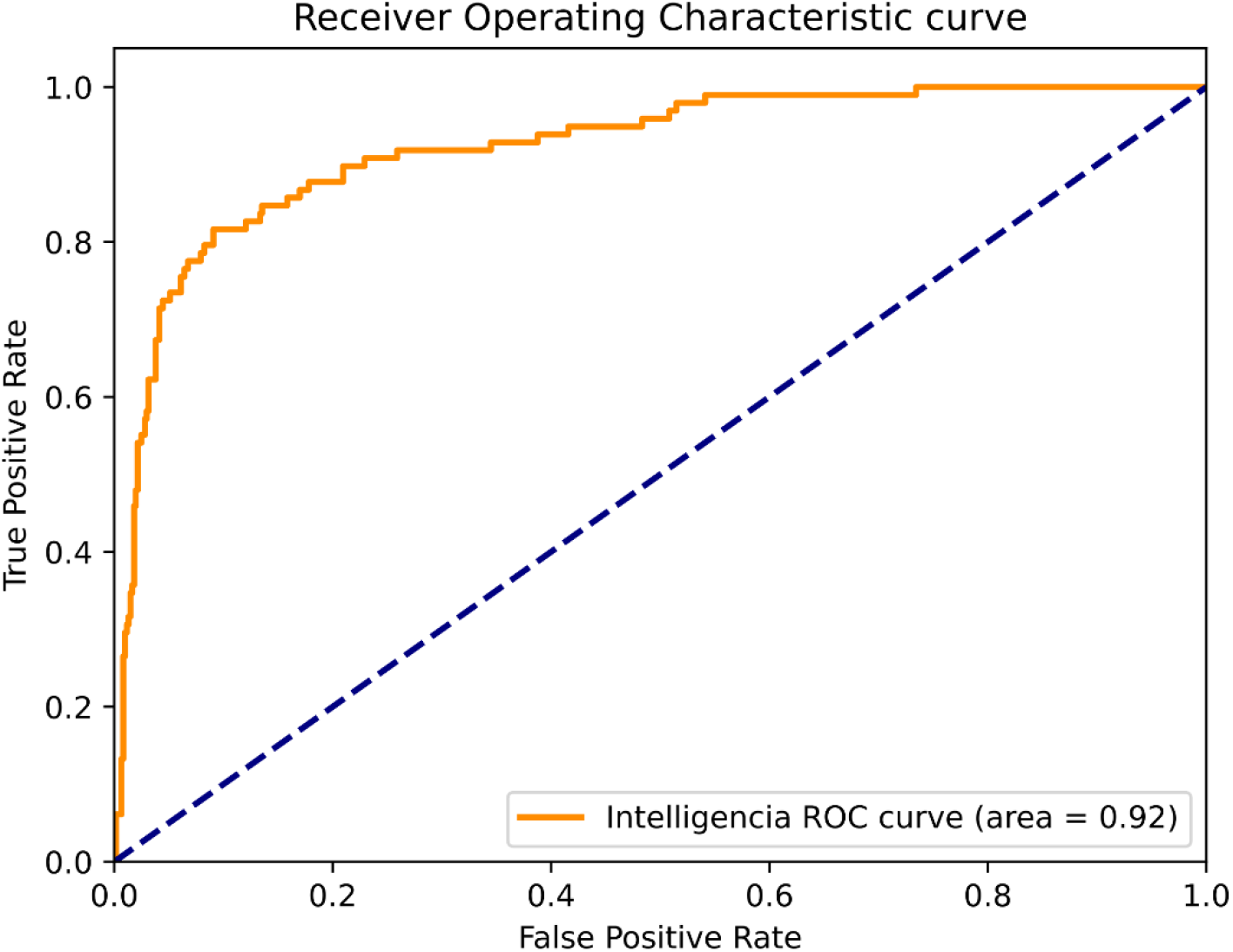
Liquid tumors at the beginning of Phase 2.

**Supplemental Figure 9:**
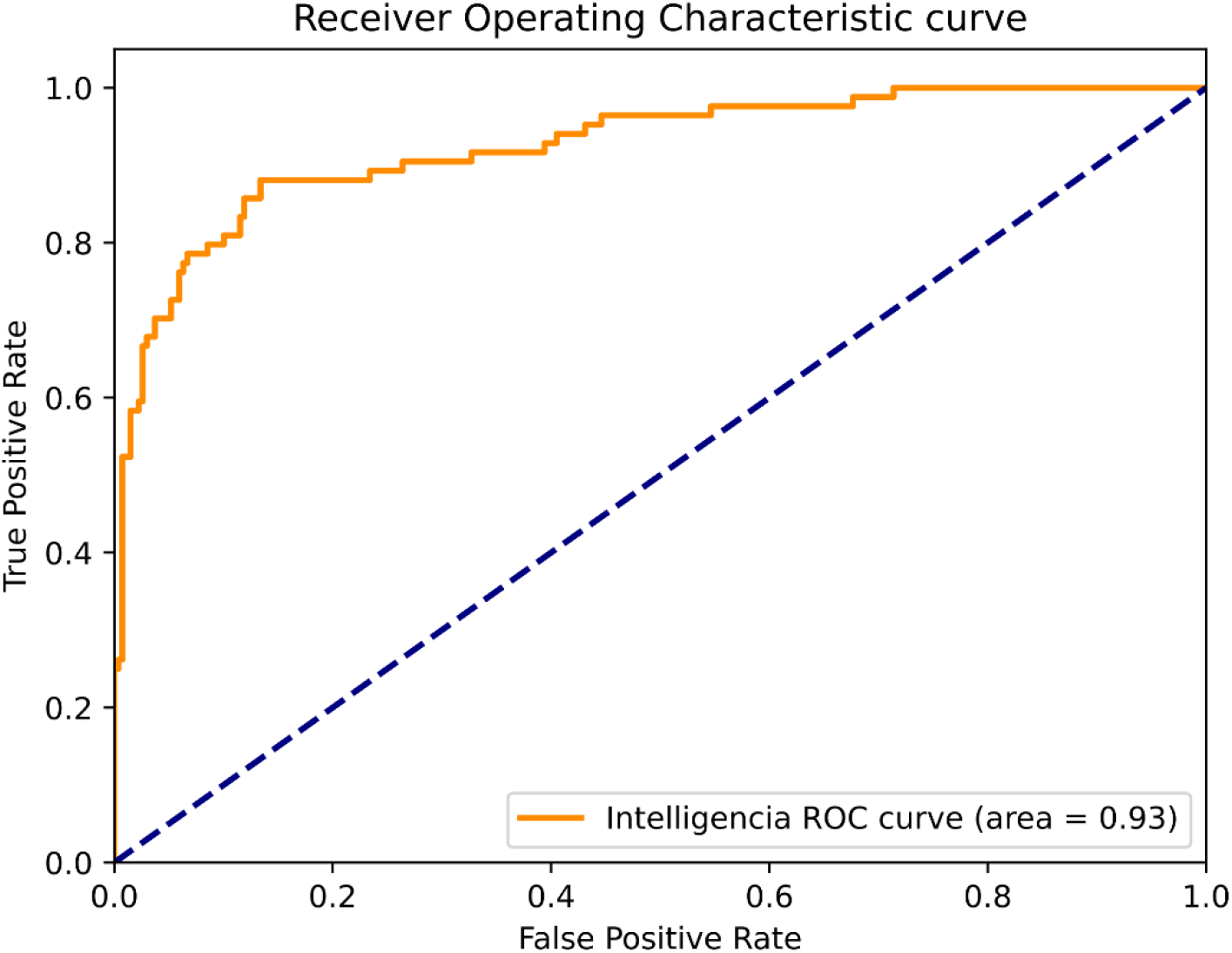
Liquid tumors at the end of Phase 2.

**Supplemental Figure 10:**
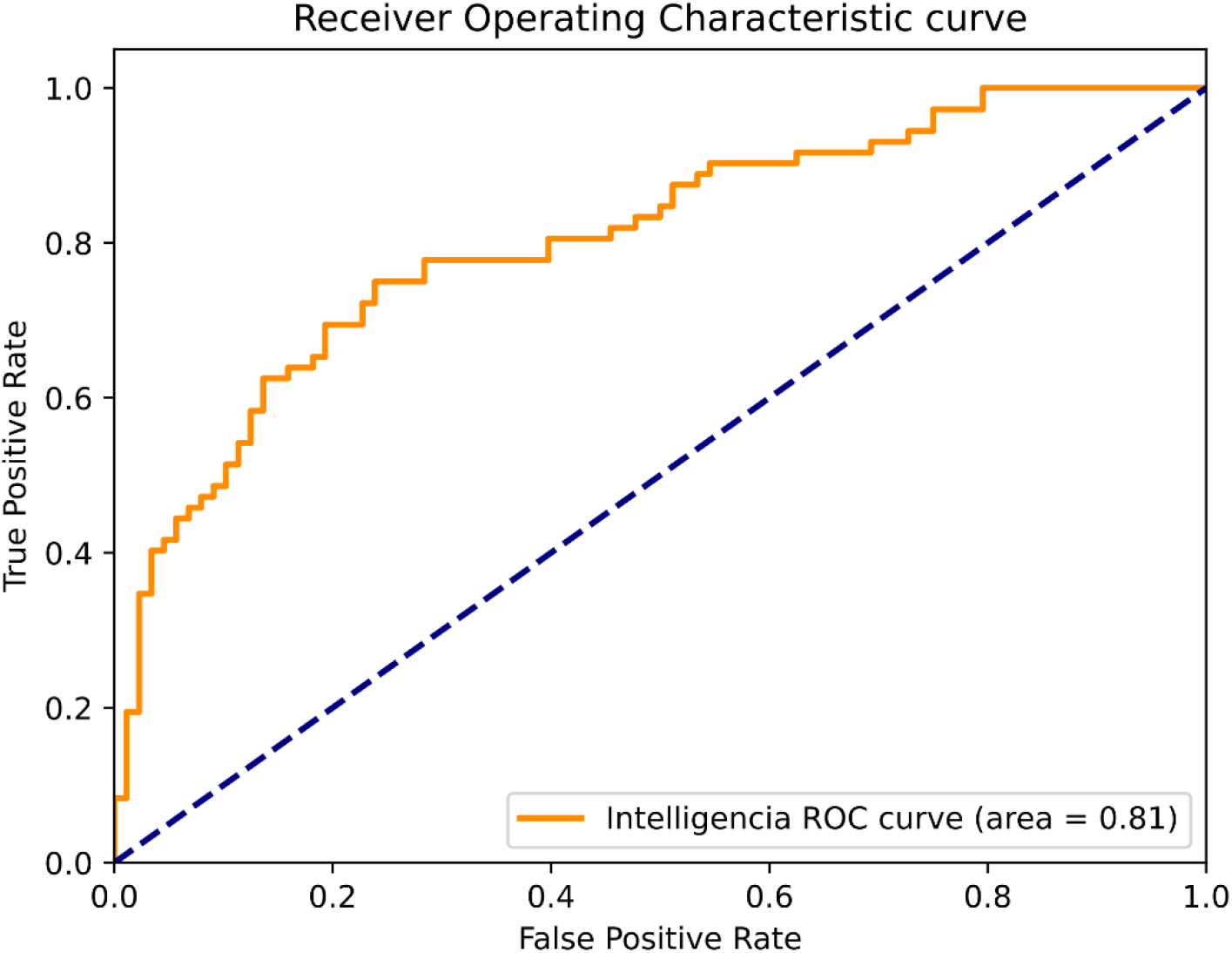
Liquid tumors at the beginning of Phase 3.

**Supplemental Figure 11:**
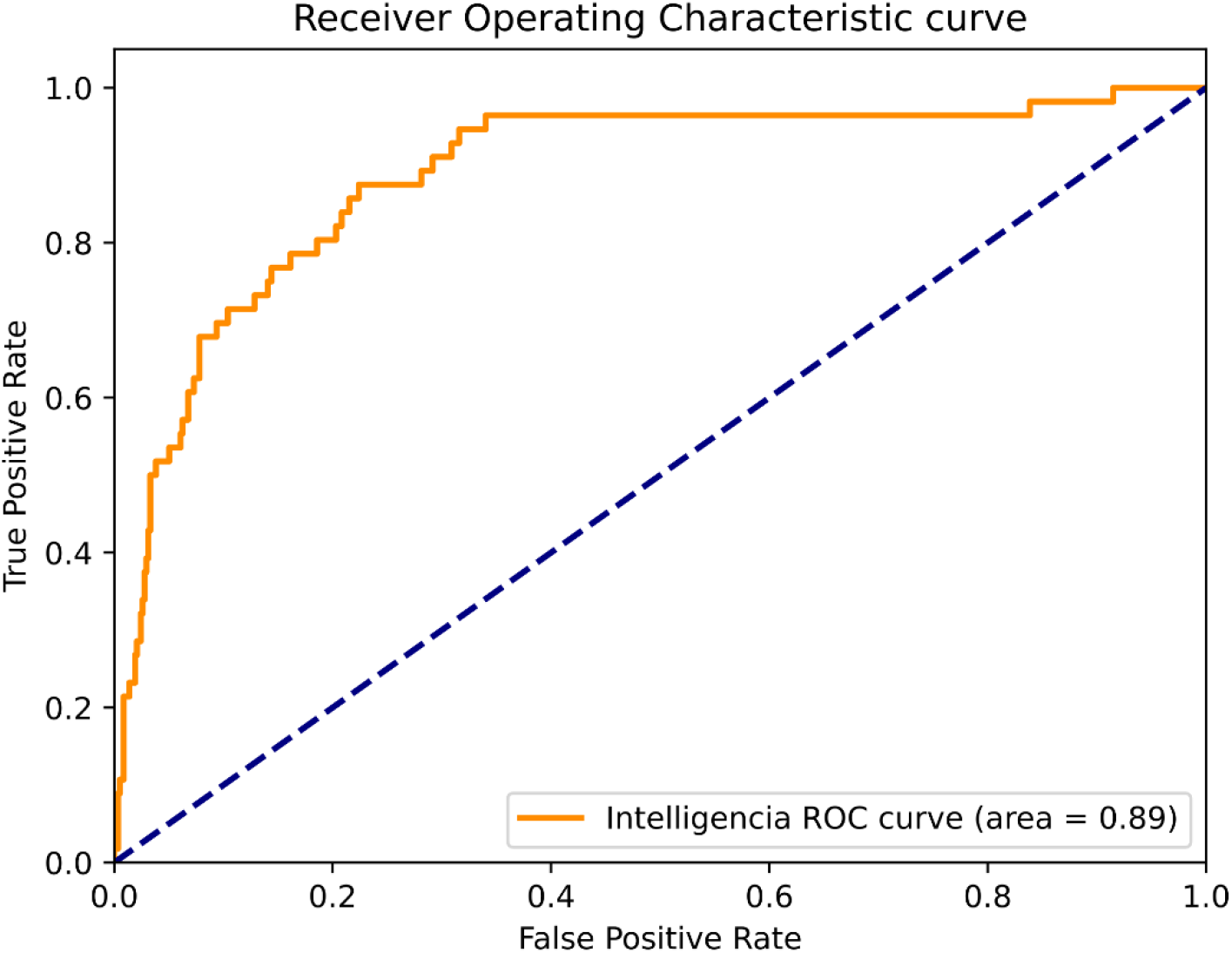
Solid tumors at the beginning of Phase 1 Post 2017.

**Supplemental Figure 12:**
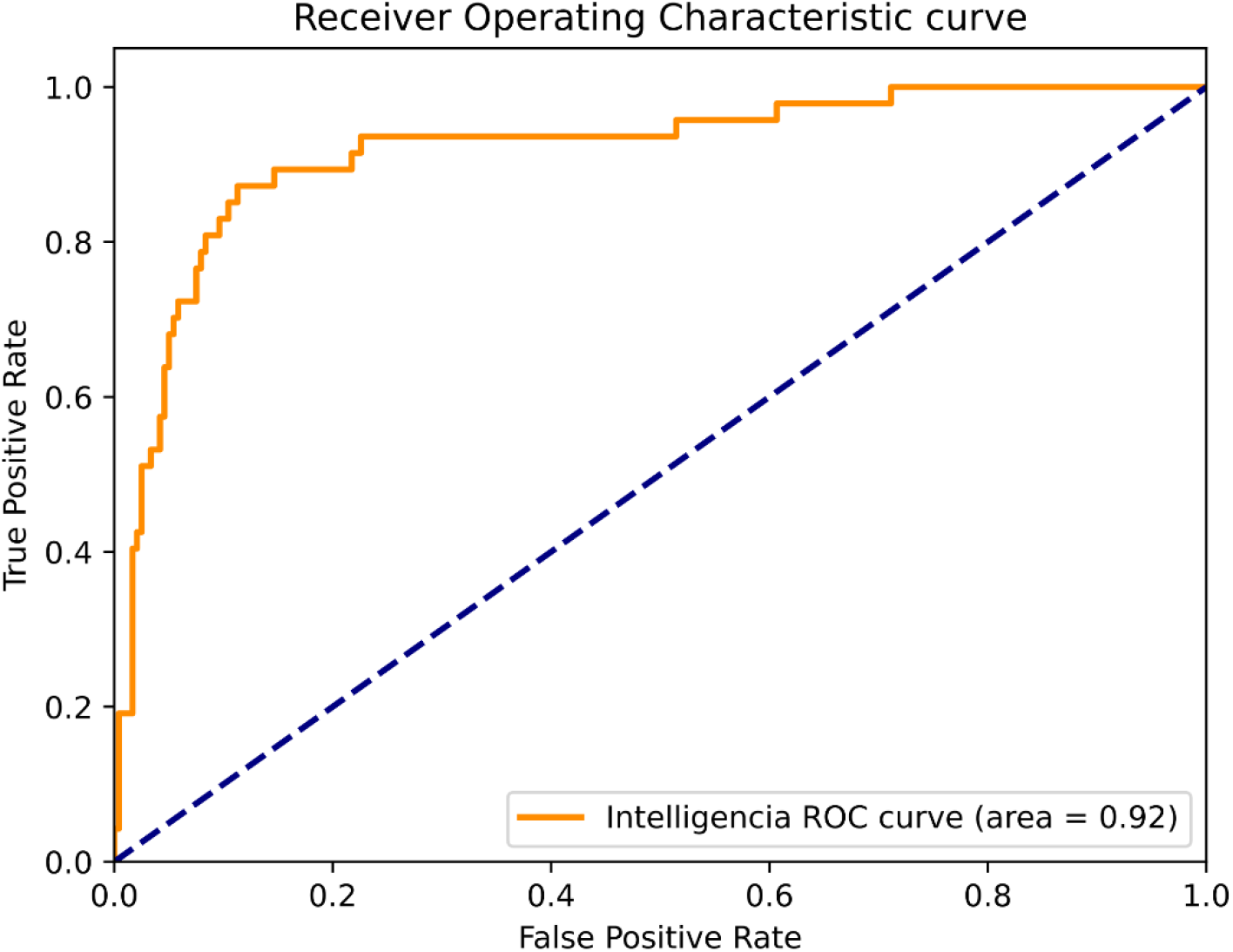
Solid tumors at the end of Phase 1 Post 2017.

**Supplemental Figure 13:**
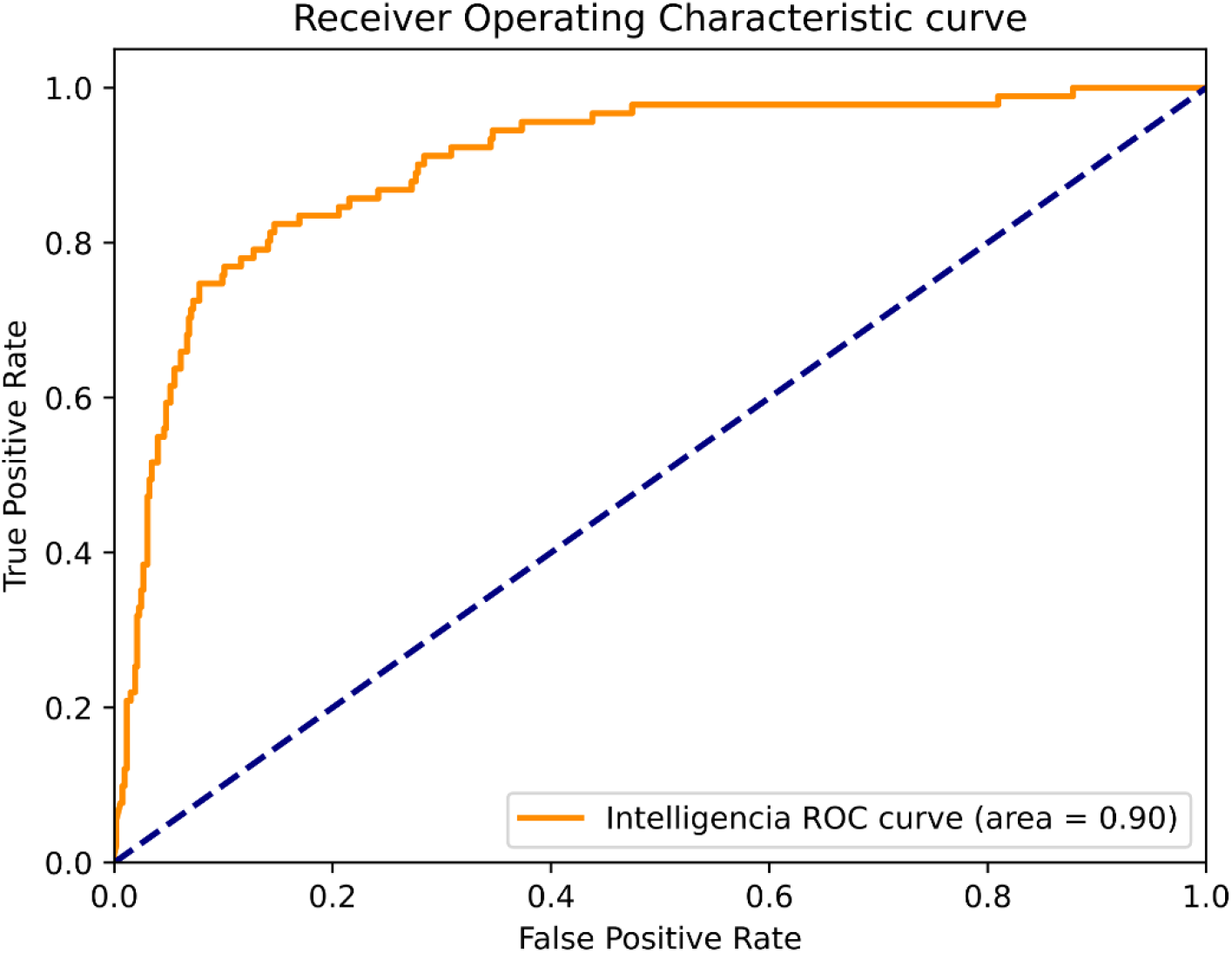
Solid tumors at the beginning of Phase 2 Post 2017.

**Supplemental Figure 14:**
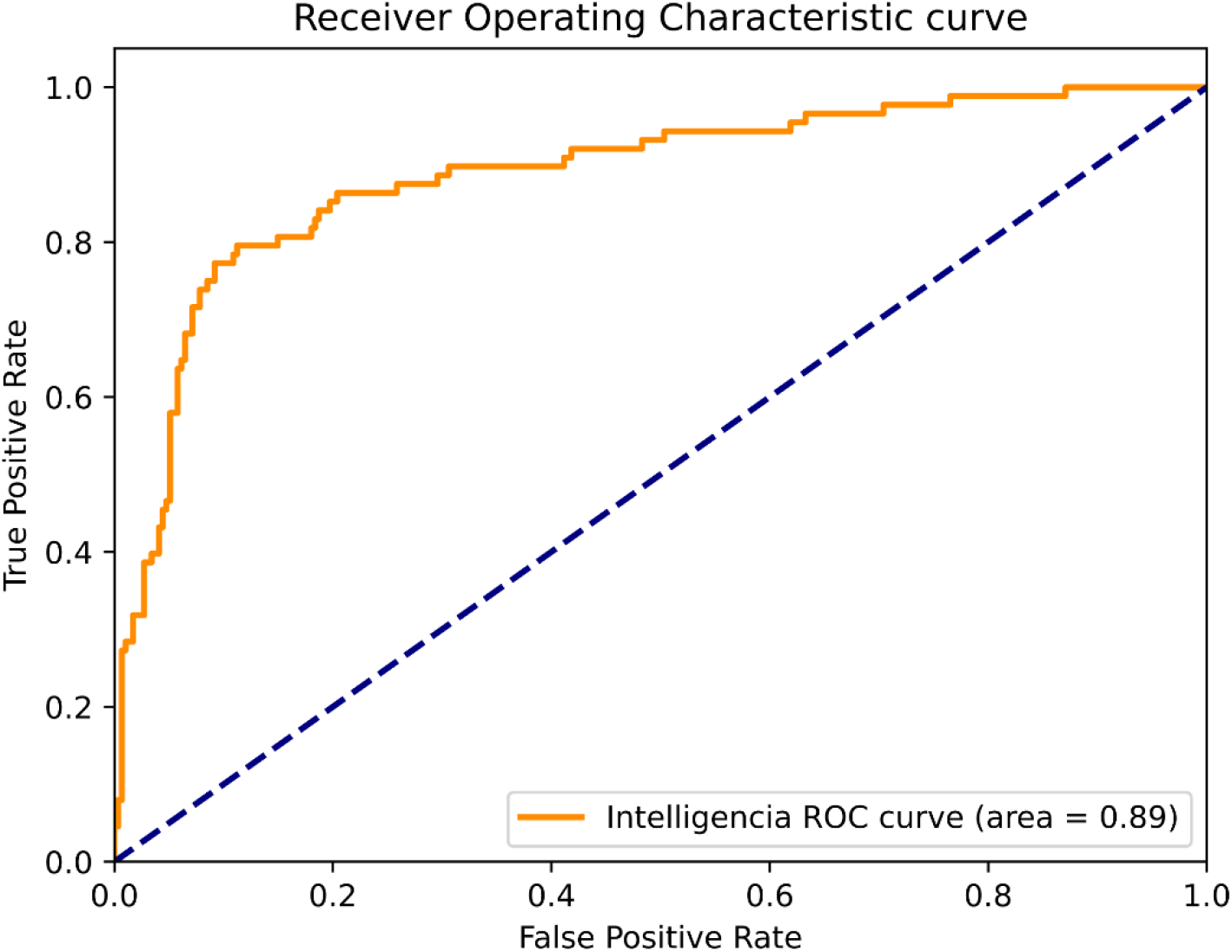
Solid tumors at the end of Phase 2 Post 2017.

**Supplemental Figure 15:**
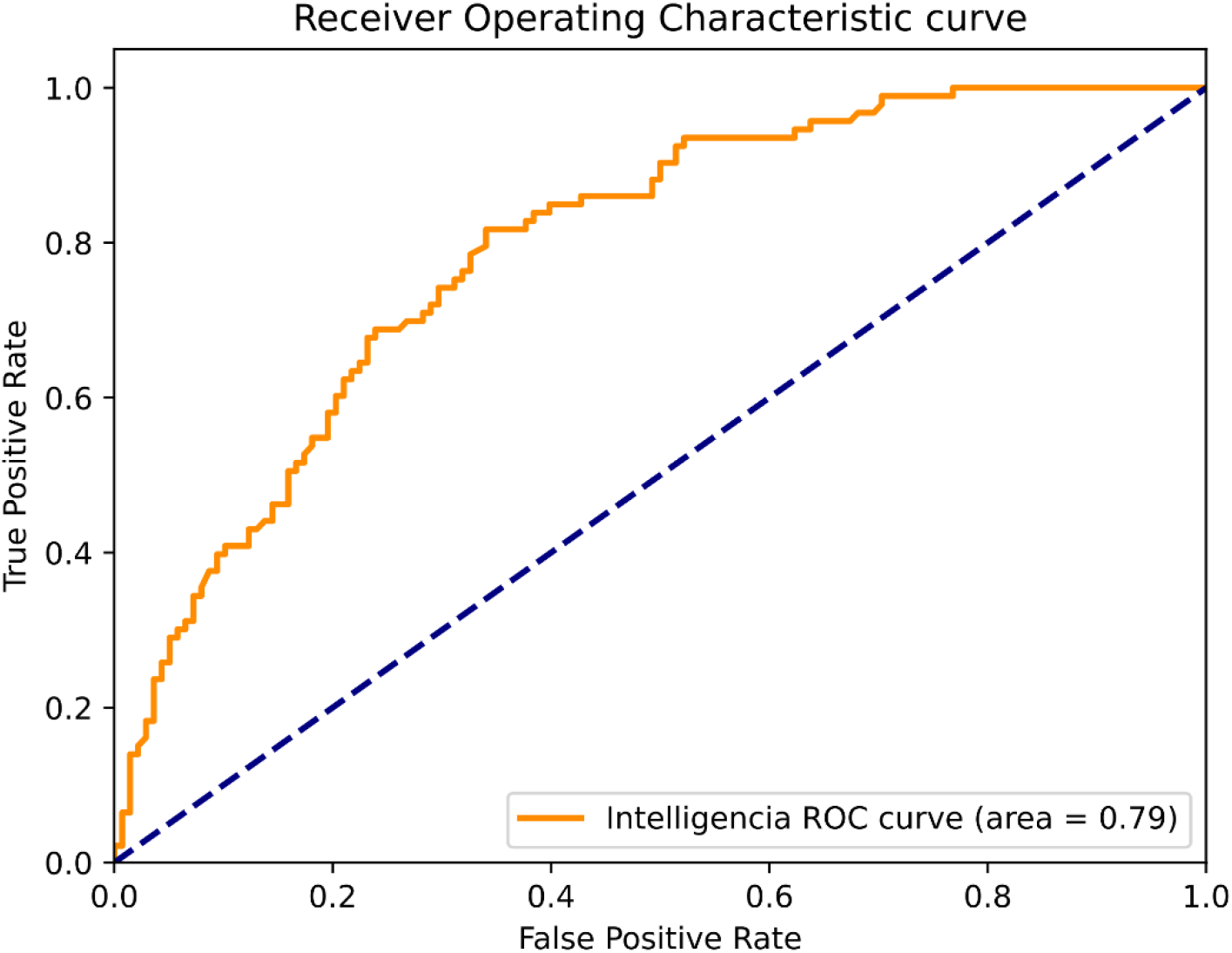
Solid tumors at the beginning of Phase 3 Post 2017.

**Supplemental Figure 16:**
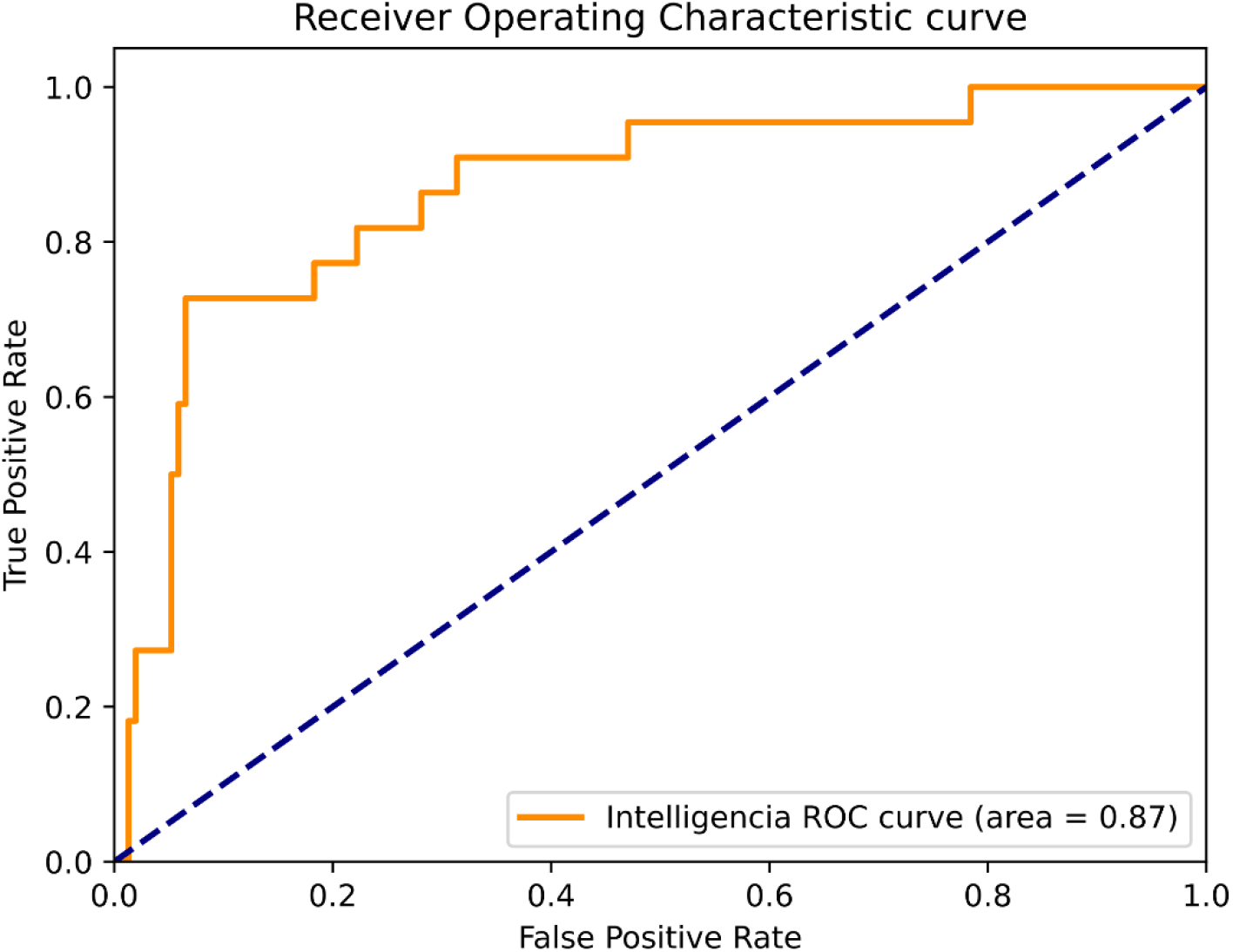
Liquid tumors at the beginning of Phase 1 Post 2017.

**Supplemental Figure 17:**
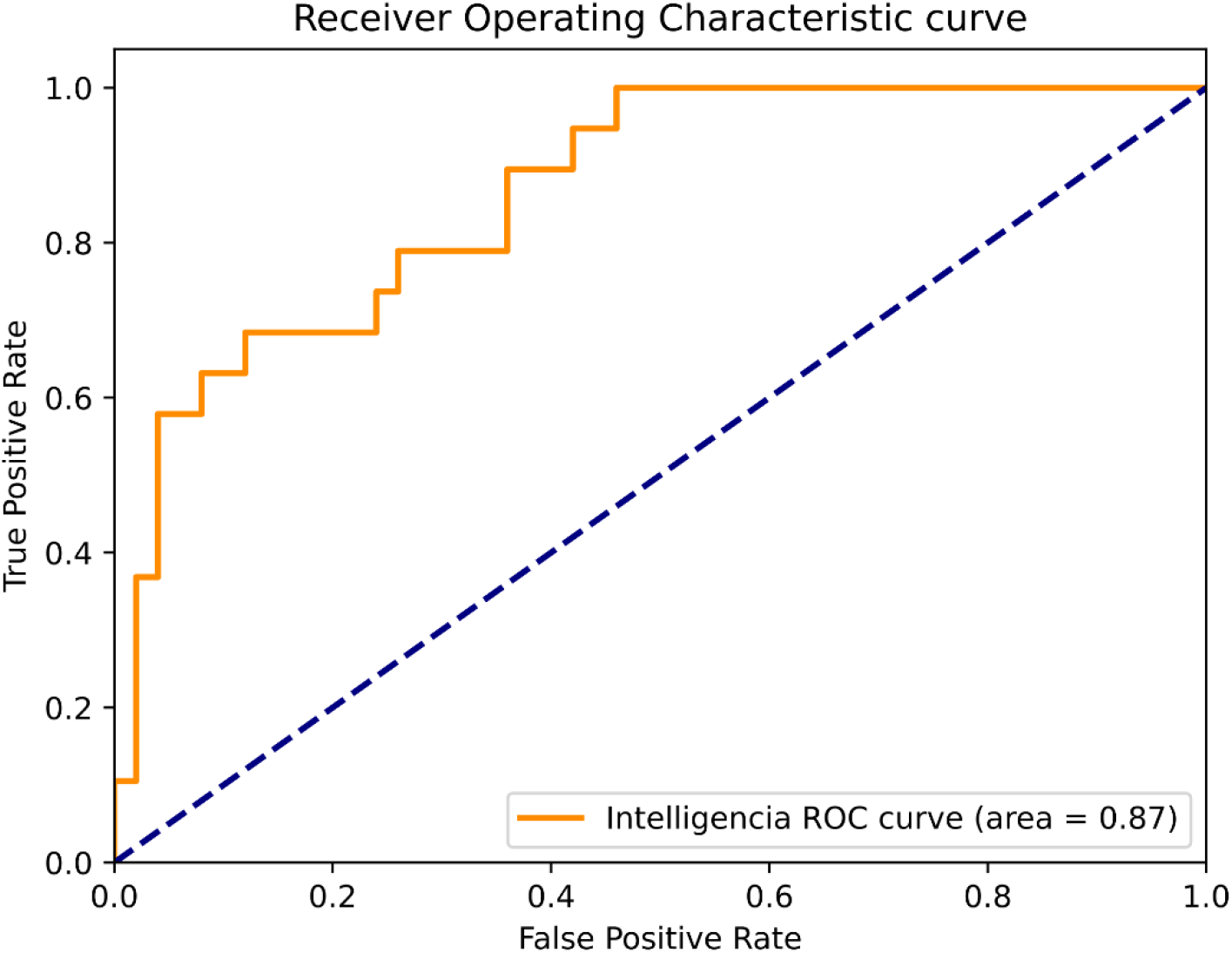
Liquid tumors at the end of Phase 1 Post 2017.

**Supplemental Figure 18:**
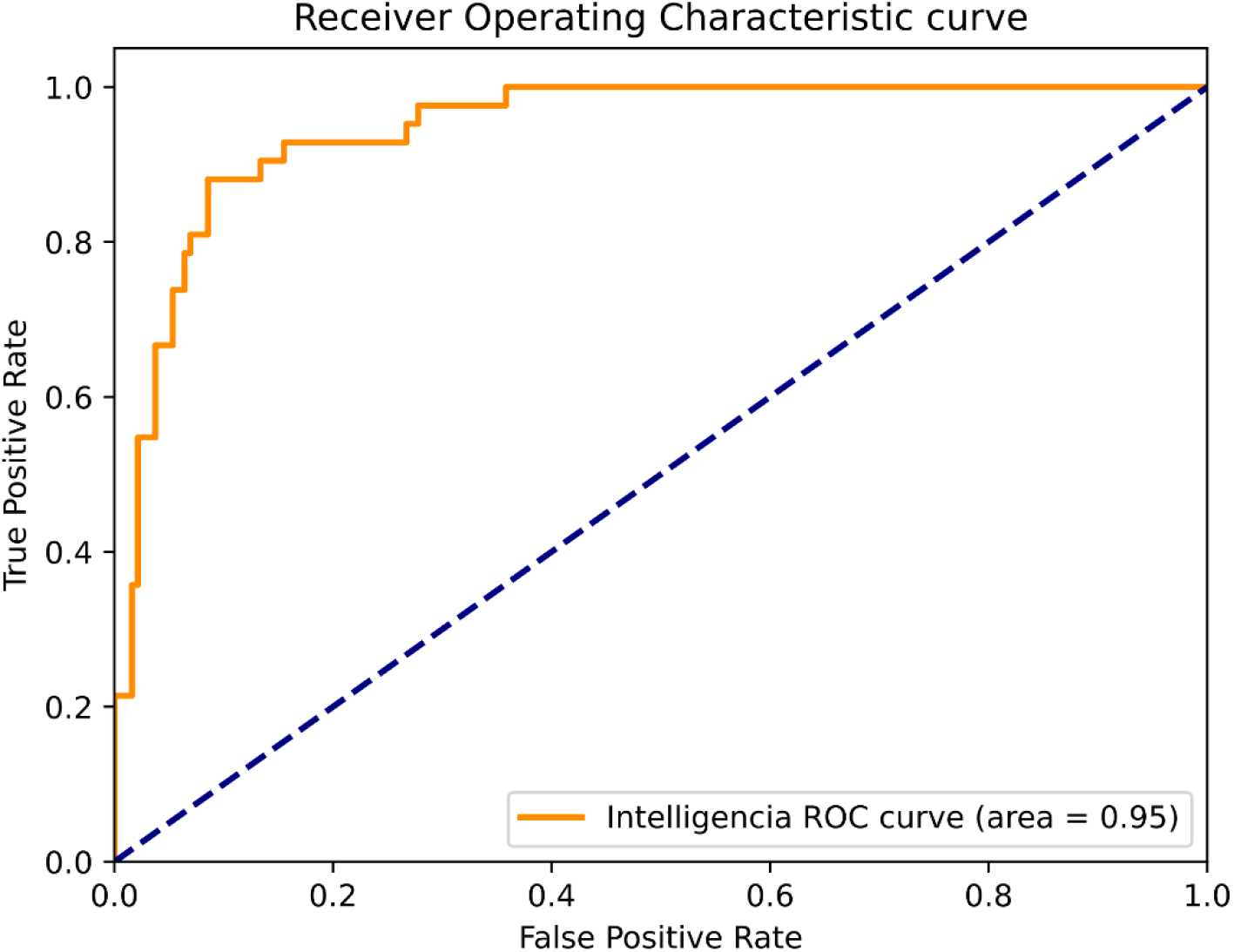
Liquid tumors at the beginning of Phase 2 Post 2017.

**Supplemental Figure 19:**
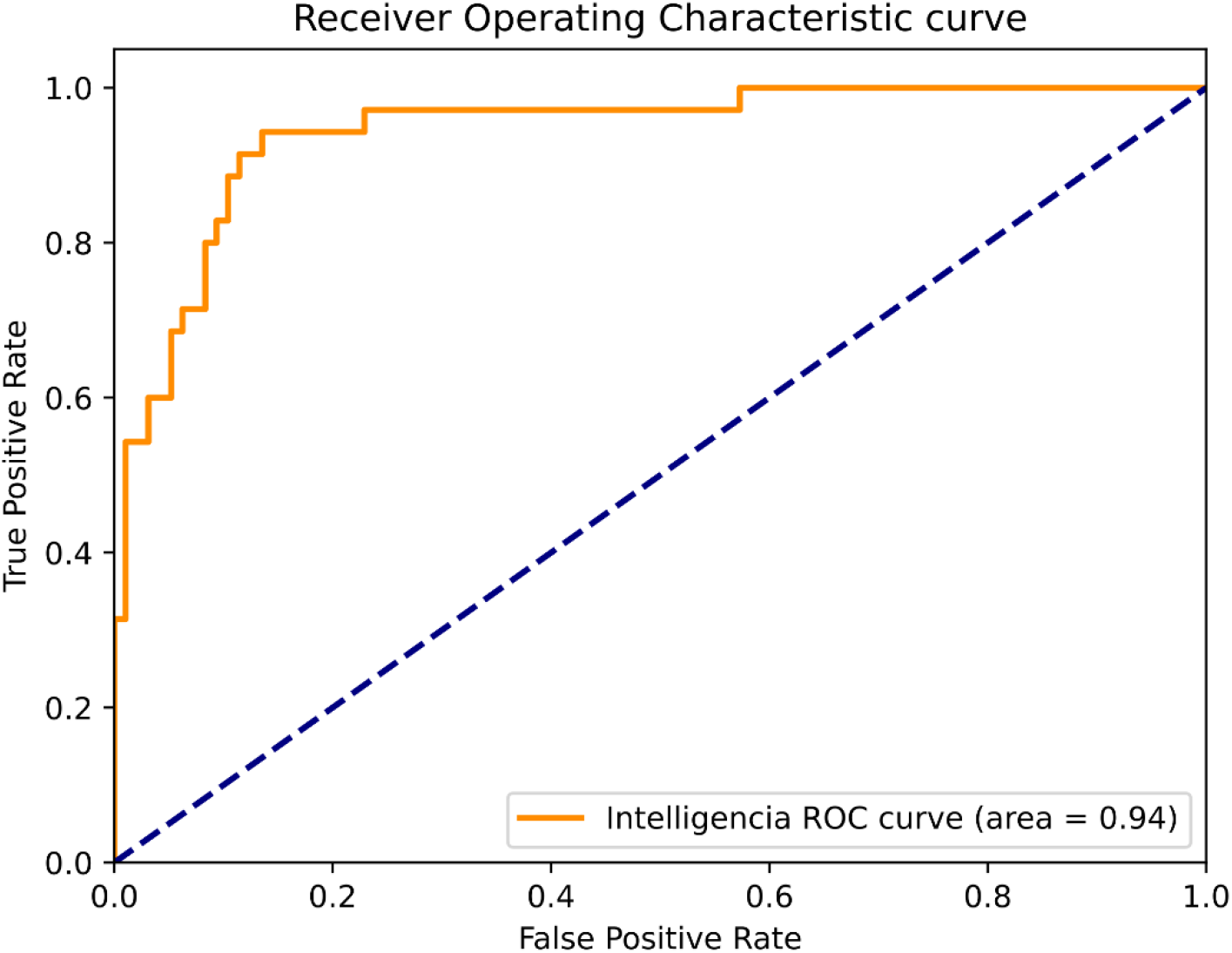
Liquid tumors at the end of Phase 2 Post 2017.

**Supplemental Figure 20:**
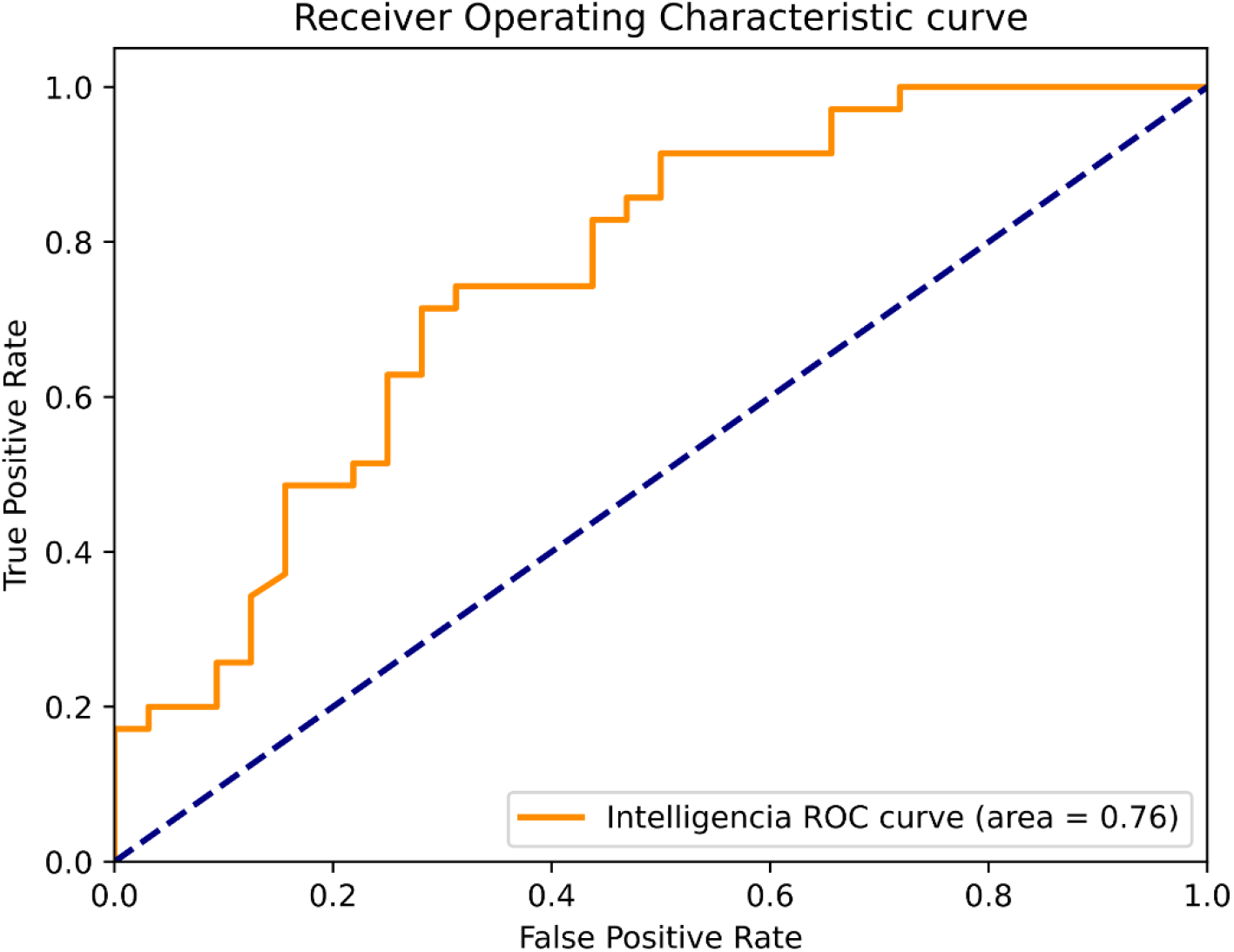
Liquid tumors at the beginning of Phase 3 Post 2017.

## Footnotes

1 We define PTRS as the probability that a development program will receive approval by the regulator (i.e., the FDA in the US). Given this, “risk” can be defined as 1-PTRS. In other words, the lower the PTRS, the higher the risk.

2 We define “development program” as the combination of a specific molecule (or combination of molecules) and a particular indication. For example, a development program can be a Phase 2 clinical trial that aims to evaluate the combination of a PD-1 molecule and a CTLA-4 molecule, for the treatment of melanoma. As a result, a particular drug (say a PD-1 inhibitor) might result in several different development programs – for example, one for say, breast cancer in Phase 1, another one for melanoma in Phase 3, etc.

3 Definition of acronyms used as example trial outcomes: ORR stands for Objective Response Rate, OS for Overall Survival, PFS for Progression-Free Survival, PSA for Prostate-Specific Antigen, CRR for Complete Remission Rate, ACR for American College of Rheumatology, MDS for Movement Disorder Society, and UPDRS for Unified Parkinson’s Disease Rating Scale.

4 Receiver Operating Characteristic

